# A monophyletic origin of domesticated barley

**DOI:** 10.1101/2025.02.26.638035

**Authors:** Cong Tan, Lingzhen Ye, Jing Chen, Chenyang Zhang, Ping Zhou, Cen Tong, Zhonghua Chen, Robbie Waugh, Chengdao Li, Tianhua He

## Abstract

The origin of domesticated barley has been debated extensively over the past century. Early botanical and comparative morphological research and recent genetic research supported a polyphyletic origin. A monophyletic origin was proposed after the discovery that a single genetic locus controls the presence of hull-less grains in all domesticated barley. Interpreting the origin of domesticated barley is further complicated by the archaeological record showing that the first domesticates had two rows of grain on the inflorescence, as does its wild progenitor. However, these two-rowed types effectively disappeared from the record for thousands of years, and they were replaced by derived six-rowed types that dominated barley’s cultivation history and only reappeared in the record around one thousand years ago. Here, we used two independent datasets with large sample sizes and genome-wide genetic markers to re-evaluate barley cultivation history. We unequivocally demonstrate that modern domesticated barley has a monophyletic origin. Phylogenetic reconstruction and examination of current archaeological records suggests that the west Fertile Crescent was most likely where barley was first domesticated. Modern two-rowed types were likely derived from six-rowed domesticated types not earlier than 3,500 years ago in modern Türkiye and Iran. We conclude that the two distinct non-brittle rachis haplotypes, coinciding with distinct row types in cultivated barley, are likely the results of two chronological events along the same evolutionary line. Our results provide a reconcilable framework for explaining genetic and phytogeographic patterns and pave the way for a consensus on the origin of barley domestication.

## Introduction

The origin of domesticated barley (*Hordeum vulgare* L.) has been constantly debated over the last century. Early botanical and comparative morphological research suggested that domesticated barley was of polyphyletic origin based on the existence of multiple centres of diversity in the geographic region of the northern Fertile and the mountain areas of Eastern Asia(Schiemann 1948). Recent genetic research favours the polyphyletic origin hypothesis, suggesting modern domesticated barley are descendants of several wild barley populations (Poets, et al. 2015; Civáñ and Brown 2017; Pankin, et al. 2018). Some researchers, however, believed that barley domestication might have been di-phyletic(Takahashi 1955; Zohary and Hopf 2000; Brantestam, et al. 2004; Saisho, et al. 2004; Kilian, et al. 2006; Tanno and Willcox 2006; Azhaguvel and Komatsuda 2007; Molina-Cano, et al. 2007). Non-brittle rachis is the primary domestication trait in barley. The wild-type genotype is *Btr*1/*Btr*2, whereas the non-brittle rachis genotype in the domesticated barley is either *btr*1/*Btr*2 or *Btr*1/*btr*2, with *btr*1/*Btr*2 (Type W) being more common among western accessions and *Btr*1/*btr*2 being dominant among eastern accessions (Takahashi 1955; Komatsuda, et al. 2004; Komatsuda, et al. 2007). The two genetic loci for non-brittle rachis are closely linked (Komatsuda, et al. 2004). Thus, the observed differences in the geographic distribution pattern of modern germplasm have been explained by two independent origins in the northern and southern Levant(Komatsuda, et al. 2004) (Azhaguvel and Komatsuda 2007). The monophyletic origin, however, cannot be easily dismissed because a single genetic locus controls the presence of hull-less grains in all domesticated barley(Taketa, et al. 2008). Meanwhile, AFLP molecular marker analysis(Badr, et al. 2000) and comparative genomic analysis of ancient barley grains(Mascher, et al. 2016) supported the monophyletic hypothesis that the Israel-Jordan area was likely the region where barley was first domesticated.

The idea of a second domestication centre on the Tibetan plateau in eastern Asia has also been a subject of debate over the past century(Aberg 1938; Schiemann 1948; Zohary 1959; Staudt 1961; Takahashi and Hayashi 1964; Morrell and Clegg 2007; Dai, et al. 2012; Pourkheirandish, et al. 2018) (Zeng, et al. 2018; Guo, et al. 2022). The finding of wild barleys with six-rowed and brittle rachises (*H. agriocrithon*) in the Tibetan plateau(Åberg 1940) was previously considered to support the hypothesis that this region was a barley domestication centre. This hypothesis was later rejected because the six-rowed wild barley was explained as feral forms contaminating cultivated varieties(Zohary 1959; Staudt 1961; Takahashi and Hayashi 1964). More recent field explorations indicated that true wild barley may be present in Tibet, Nepal, India, Pakistan, and Afghanistan(Yang and Yen 1985; Shao and Li 1987; Corke and Atsmon 1990). Pourkheirandish recently discovered that some *H. agriocrithon* barley accessions carried *Btr1* and *Btr2* haplotypes not found in any cultivars and argued that the occurrence of six-rowed wild types implies the six-rowed allele pre-dating domestication, thus favouring the hypothesis that six-rowed cultivated barley was derived from *H. agriocrithon*(Aberg 1938; Pourkheirandish, et al. 2018). However, Guo *et al*. used genome-wide marker data and whole-genome resequencing to show that all *H. agriocrithon* accessions of a germplasm collection are hybrid forms that arose by admixture of diverse domesticated and wild populations(Guo, et al. 2022).

Current archaeological evidence does not dispute that the earliest domesticated barley is dated to ≈8,500 cal. BC. and was the two-row type (Van Zeist 1970; Hillman, et al. 1989). Six-rowed barley split off from its two-rowed parent shortly after cultivation began and is expressed as a fully domesticated crop plant in the archaeological record (Rollefson and Simmons 1985; Willcox 1998). Six-rowed barley dominated at the Balkans, Central Europe, Southern Europe, and North Africa 7,000–4,000 years before the present (BP) and only six-rowed barleys were cultivated in the vast agricultural area east of the Hindu Kush until a hundred years ago (Clark 1967). After it disappeared from ancient Mesopotamia before the end of the fifth millennium BC, cultivated two-rowed barley only reappeared *c.* 700 BC in the archaeological record of Beycesultan in Anatolia (Helbaek 1961), leaving a gap of over three thousand years separating the two-rowed barleys of Beycesultan from those of the later Halafian period (c. 5,400–5,000 BC) and no authentic records of cultivation of two-rowed type bridging the gap. Thus, one more question was asked more than half a century ago(Clark 1967) and remains to be answered. Does two-rowed barley have a continuous but un-recorded history in the area adjacent to where it was initially domesticated, or must the two-rowed barleys be a second and independent introduction of wild barley into cultivation?

The archaeological sequence of barley domestication events seems reliable and less disputable (Clark 1967; Fuller, et al. 2011), while the botanical and genetic explanations need more consensus. Advancements in genetic technology and statistical approaches allow for inference of the phylogenetic pattern of evolutionary history, thus offering answers as to whether a crop was monophyletic or polyphyletic (Badr, et al. 2000; Morrell and Clegg 2007; Dai, et al. 2012; Poets, et al. 2015; Civáñ and Brown 2017; Pankin, et al. 2018; Zeng, et al. 2018). However, modern germplasm collections are incomplete due to climatic changes and human impacts over millennia (Fuller, et al. 2011), which was made worse by often a small number of accessions being investigated in previous studies. Meanwhile, previous analysis of evolutionary history using a neighbouring joining approach or principal components clustering offers no consideration for local extinction, which further fuelled the dispute on the origin of domesticated barley (Fuller, et al. 2011; Allaby, et al. 2017).

Here, we conduct analyses on the origin and evolution of domesticated barley using two independent datasets. We first analysed 340 worldwide accessions with publicly available high-coverage whole-genome sequences, including wild barley and landrace from Tibet, with wild barley from Israel as the reference genome for variant discovery. We re-examined a dataset of 5,448 worldwide wild and landrace barley accessions with genome-wide SNP profiles. The Genebank of the Leibniz Institute of Plant Genetics and Crop Plant Research (Gatersleben, Germany) collected those barley accessions from all countries and regions with barley cultivation over the past century and genotyped them recently (Milner, et al. 2019). Thus, our datasets include a substantial expansion of the sampling of wild accessions and landrace throughout the geographic range of barley (**Figure 1 and Supplementary Table 1**). We used time-calibrated phylogenies to reconstruct the evolutionary history of domesticated barley, which considered historical speciation and extinction rates by fitting stochastic birth-death models through Markov chain Monte Carlo simulations (Louca and Pennell 2021). We also sieved the archaeological records to trace the early domestication of barley in the Fertile Crescent and adjacent regions. Collectively, our findings support a monophyletic origin of domesticated barley. We suggest that six-rowed wild barleys in Tibet are likely the relics of the early introduction of six-rowed barley into Tibet *c*. 7,300 years ago and rewilded. We conclude that the original domesticated two-rowed barley may be extinct, and the modern two-rowed type was derived from the six-rowed types on multiple occasions thousand years after the two-rowed barley was first domesticated and became extinct.

**Figure 1.**
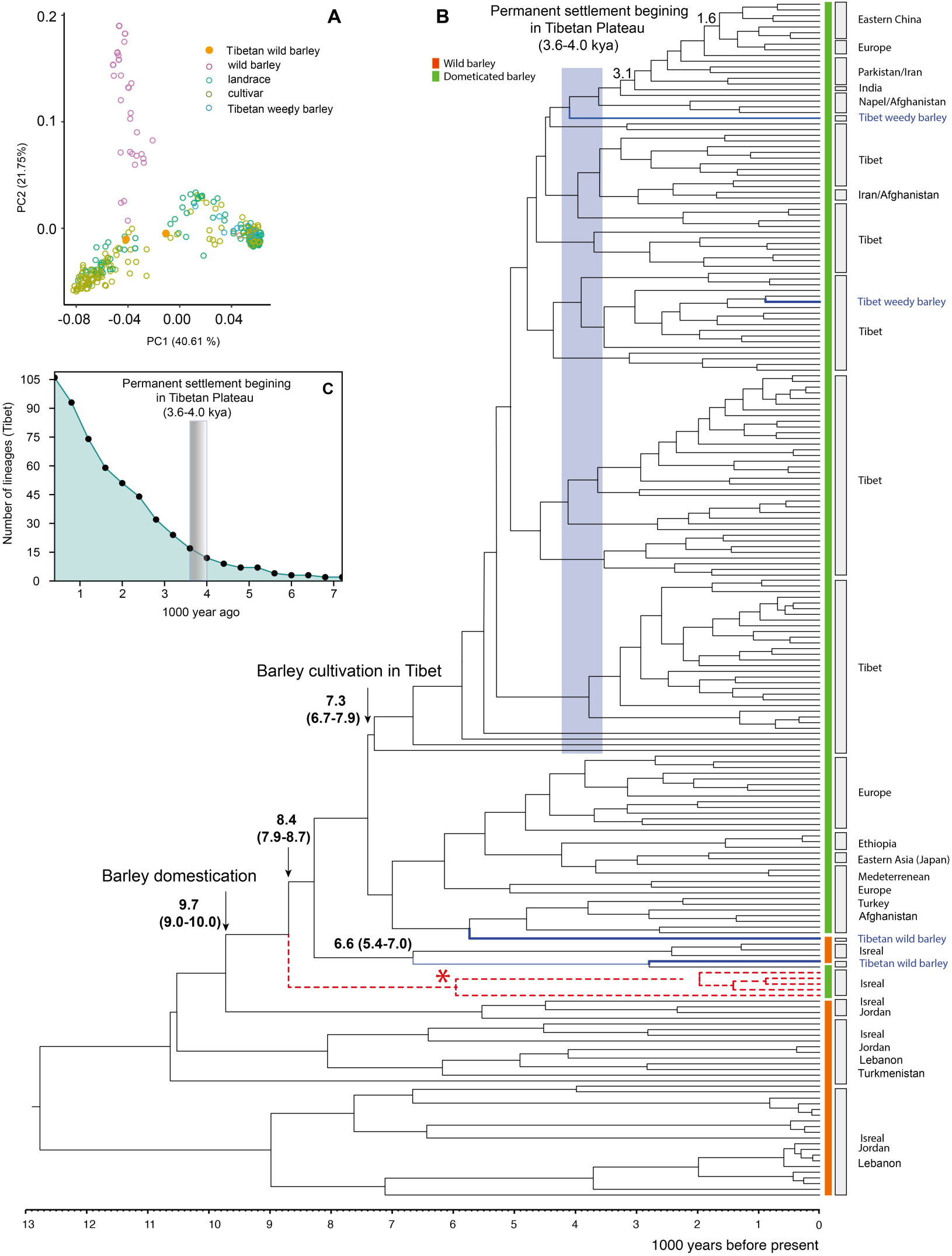
Origin of domesticated barley. **A**, Principal component analysis of 340 barley accessions; **B**, Time-based phylogenetic analysis on 340 barley accessions with deep coverage sequences and wild barley genome as reference. The dotted lines represent the ancient barleys discovered from Yoram Cave in the Judean Desert, used as the molecular clock calibration point. **C**, The establishment of permanent settlement in the Tibetan Plateau.

## Materials and Methods

### Origin and Evolution analysis of modern domesticated barley

We used two independent datasets to explore modern domesticated barley’s origin and evolution. We searched the National Center for Biotechnology Information (NCBI) and European Nucleotide Archive (ENA) for barley whole-genome resequencing data. We obtained 340 barley accessions with >5× coverage whole-genome shotgun resequencing data (**Supplementary Table 1**). Those barley accessions included wild barley (54), landraces (155) and cultivars (131) from 38 countries/regions. Two wild barley accessions and semiwild barley (also known as weedy barley (Zeng, et al. 2018)) from Tibet in China were included. We also obtained whole-genome sequencing data for five ancient barley grain samples discovered from Yoram Cave in the Judean Desert in Israel, dated 6,000 years ago(Mascher, et al. 2016). The sequences were mapped to Israel’s wild barley reference genome EC_S1 (Pan, et al. 2023; Zhang, et al. 2023), and whole genome SNP variants were identified using the Sentieon mapping and variant calling pipelines (Freed, et al. 2017), respectively. We filtered out variants with a missing rate of over 10% and a minor allele frequency (MAF) of less than 0.05. Principal component analysis was conducted by PLINK v2.0 (Chang, et al. 2015) to reveal the visual genetic relationship between groups of barley germplasms.

Phylogenetic relationships among the 340 barley accessions were explored with Bayesian Markov chain Monte Carlo under GTR substitution and Yule process evolution models using BEAST v1.10.4 (Suchard, et al. 2018). We employed an uncorrelated relaxed molecular clock with one calibration point to create the time-based phylogenetic tree. We set the five ancient barley samples as a monophyletic group, and the crown age of this group was set at 6,086 cal. BP(Mascher, et al. 2016) as the calibration point. We used the lognormal distribution function as priors, as this distribution allows the assignment of the highest point probability for the node age that must be older than the designated age at the calibration point. We ran analyses of 100 million generations, sampling every 2,000 generations. We accepted that the first 10 million generations from the Markov chain Monte Carlo (MCMC) sample were a conservative burn-in. We carried out five independent Monte Carlo Markov chain runs, and the trees were combined with LogCombiner (Suchard, et al. 2018), and a maximum clade credibility tree was then generated with TreeAnnotator (Suchard, et al. 2018). Final trees were inspected and edited with FigTree v. 1.4.4 (http://tree.bio.ed.ac.uk/software/figtree/).

### Large barley diversity panel dataset

We used a large barley diversity panel dataset to explore the origin and evolution of the two-rowed and six-rowed domesticated barley. Ready-to-use phenotypic data, including row type and region of origin, and genome-wide SNP dataset for approximately 19,778 barley (*H. vulgare* L.) accessions were made available by the Federal Ex situ Gene Bank for Agricultural and Horticultural Plant Species (IPK) in Germany. The panel includes domesticated barley (cultivars and landraces) and its conspecific wild progenitor, *H. vulgare* ssp*. spontaneum* (K. Koch). The row type was assessed during seed regeneration using plots of at least 3 m^2^ (Milner, et al. 2019). SNP profiles were derived from a single plant of the accessions in the IPK barley collection through the genotyping-by-sequencing method (Milner, et al. 2019). We retained samples with phenotypic (row type and region of origin) and genotypic data for our analysis. We kept samples designated as “landrace” and “wild barley” by excluding cultivars in our analysis, as artificial crossing may be involved in cultivars, thus confusing the phylogenetic reconstruction. The retained samples were further filtered with all samples with <5% missing genotypes and a minor allele frequency of >0.05. Consequently, we have obtained 5,802 polymorphic SNPs for 5,448 samples, including 419 wild types, 1,423 two-rowed and 3,606 six-rowed landraces, from all countries with traditional barley cultivation (**Supplementary Table 2**).

With the SNP profile of the 5,449 samples, we first used RAxML-NG to construct the phylogenetic tree following a maximum likelihood procedure (Kozlov, et al. 2019). The phylogenetic tree was then dated with MEGA 11 (Tamura, et al. 2021). We set the boundary of the crown age of all domesticated barley at 10,100 cal. BP (the results from the above analysis with 340 barley samples with deep sequencing). We trace the trait along the phylogenetic tree to explore the potential differentiated origin of the two-rowed and six-rowed barley. If the two sister accessions/lineages have the same row type, their common ancestor is assigned the same row type. The assignment process was continued till the two lineages had different row types. Meanwhile, the sequence of known haplotypes of the vrs1 gene was obtained from Saisho, et al. (2009), and the phylogenetic relationship of the haplotypes was reconstructed with RAxML-NG (Kozlov, et al. 2019). To explain the origin of domesticated two-rowed and six-rowed barley, we exhaustedly searched available literature about archaeological records of barley domestication. We started with several key papers (Clark 1967; Van Zeist and Bakker-Heeres 1982; Meadows 2005; Riehl 2019) and searched the references these key papers cited and references that cited these papers. We limited our search to records that reported the row type of the barley remains. We noted the archaeological sites and the date. The carbon dating time was converted to calibrated BC for consistency (Reimer, et al. 2020).

## Results

We collated sequence data for 340 worldwide barley accessions from NCBI and ENA, including 54 wild barleys from Israel, Jordan, Lebanon, Turkmenistan, and Azerbaijan, 288 landrace and cultivars from 38 countries (**Supplementary Table** 1). Using a wild barley genome from Israel as the reference genome, we discovered 28,788,523 high-quality SNPs and 1,917,494 InDels. Principal component analysis showed that all wild barley, except the two wild barley candidates from Tibet, formed an independent group and separated from domesticated barley accessions, suggesting a clear genetic distinct between wild and domesticated barley (**Figure 1A**).

Phylogenetic analysis using the Bayesian MCMC method revealed that modern domesticated barley is a monophyletic group (**Figure 1B**). The monophyletic domesticated barley separated from wild barley in Israel 8.4 thousand years ago (kya) (7.9-8.7 kya, 95% highest posterior density, HPD). The ancient domesticated two-rowed barley discovered from Yoram Cave in Israel (Mascher, et al. 2016) was basal to all modern domesticated barley and their direct wild ancestors from Israel. The monophyletic group, including the ancient domesticated two-rowed barley and all modern barley, is directly derived from wild barley in Israel-Jordan 9.7 kya (9.0-10.0 kya, 95% HPD). Landraces and cultivars in modern Tibet formed a monophyletic group from 7.3 kya (6.7-7.9 kya, 95% HPD). Barley accessions in Tibet diversified *c.* 4.2 kya (Fig 1 C), slightly earlier than the time with the establishment of permanent settlement in the Tibetan Plateau 4.0-3.6 kya (Chen, et al. 2015).

To further clarify the origin of domesticated barley and the different opinions centred on the origin of two-rowed and six-rowed barley, we examined a large dataset containing 5,449 diverse barley samples (Milner, et al. 2019) from 22,626 barley accessions genotyped using a GBS method, including wild barley, landraces, and cultivars, collected over the past 100 years worldwide with barley cultivation. Along with data on genotype and geographic origin, they also reported the row types for a proportion of samples. With the filtering criteria, we consequently examined 5,448 samples, including 419 wild types and 5,030 landraces comprising 1,856 two-rowed and 3,593 six-rowed types (**Figure 2, Supplementary Table** 2), with 5,802 polymorphic SNPs. Principal component analysis revealed the 5,449 samples formed four major groups with some overlapping (**Figure** 2). Wild barley formed a genetic cluster overlapping with six-rowed barley that separated into two clusters, an Asia cluster and a European cluster. Six-rowed barley from Iran, Iraq, Afghanistan, Libya and Türkiye showed a closer genetic relationship with wild barley. It sat between Asia and European clusters in the PCA plot (**Figure** 2). Two-rowed barley clustered together with some overlapping with six-rowed barley but not with wild barley clusters, suggesting a more distant genetic relationship of modern two-rowed barley to wild barley than to the modern six-rowed barley.

**Figure 2.**
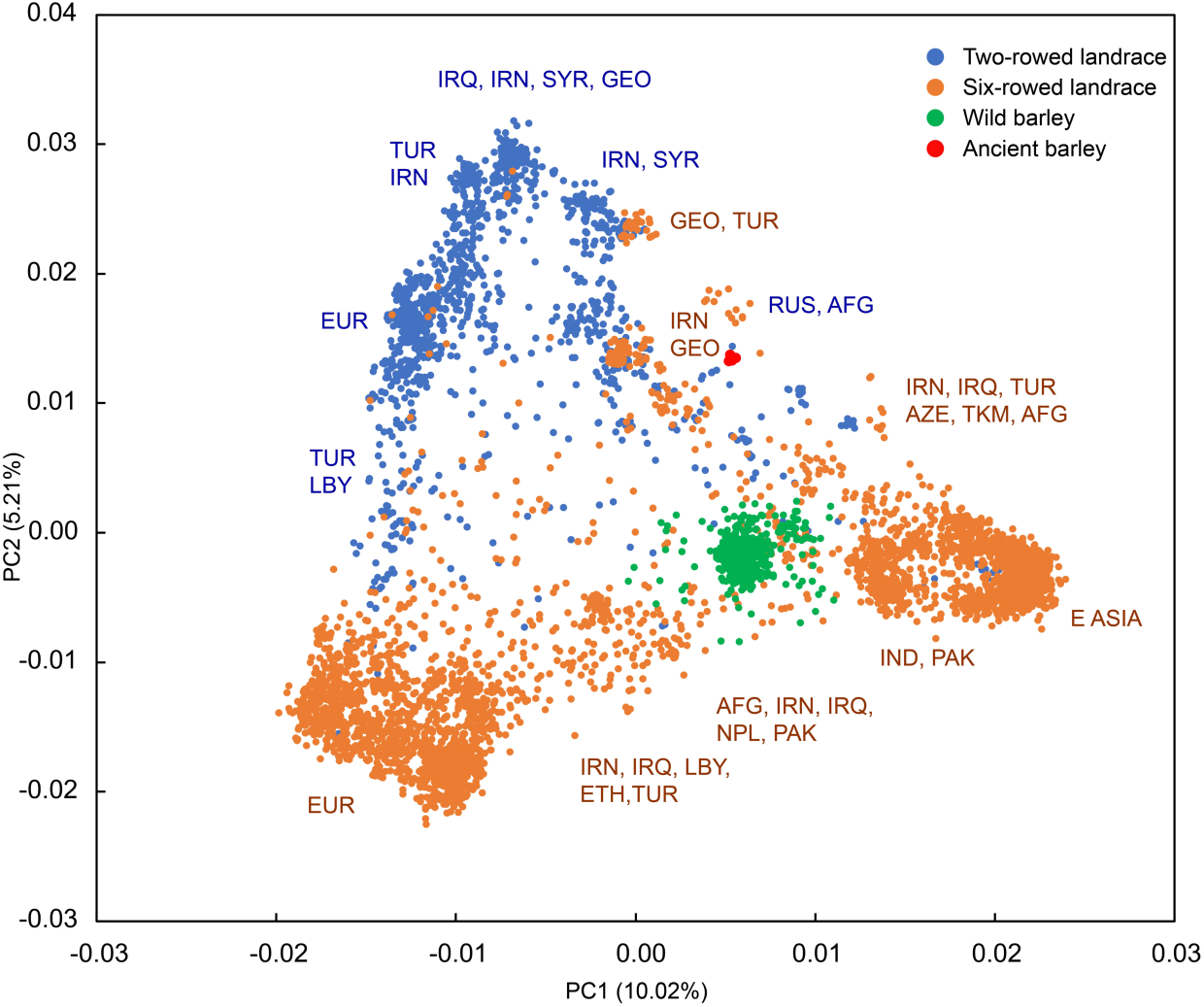
Principal component analysis of the 5,448 wild barley and landraces samples. The PCA plots showing that modern two-rowed barleys have a more distant genetic relationship with wild barley than modern six-rowed barley. Abbreviations for countries and regions were explained in **Supplementary Table** 3.

Maximum likelihood phylogenetic reconstruction with the 5,449 samples and 5,802 polymorphic SNPs showed that all 5,030 modern barley landraces, including two-rowed and six-rowed types, formed a monophyletic group, suggesting a monophyletic origin of modern barley (**Figure 3**A). Modern barley showed a closer phylogenetic relationship with wild barley from Israel than those from any other regions. The monophyletic modern barley separated from wild barley c. 13,740 cal B.P. (ya). However, ancient trait reconstruction suggests that the common ancestor of modern domesticated barley was six-rowed and started diversifying *c*. 10,400 ya. Two major groups of six-rowed barley formed from *c.* 9,000 ya., with one cluster containing six-rowed barley from Türkiye, Libya, and centre Asia, and Europe, while another cluster containing six-rowed barley from Iran, east of Hindu-Kush, and eastern Asia. Small clusters of six-rowed barley also existed in The Fertile Crescent and adjacent regions, i.e. Iraq, Iran and Syria (**Figure 3**A).

**Figure 3.**
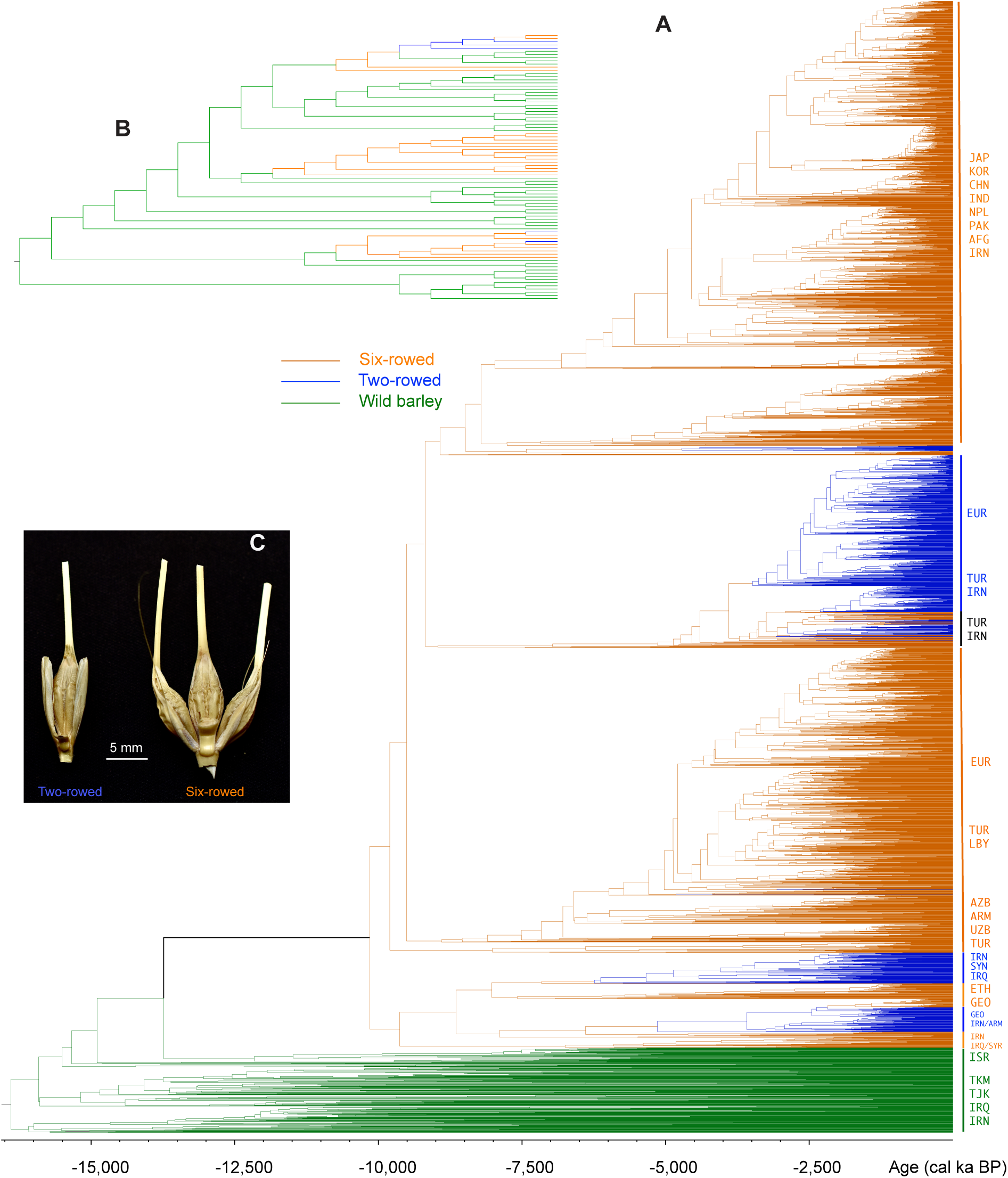
Domestication and evolution of two-rowed and six-rowed barley. **A**, Phylogeny of the 5,449 samples built from 5,802 polymorphic SNPs. **B**, Phylogeny (not time-based) of the *vrs1* gene haplotypes corresponding to row types. **C**, Grains of two-rowed barley and six-rowed barley. Abbreviations of country and region were explained in **Supplementary Table** 3.

All modern two-rowed barleys were derived from six-rowed in the phylogeny (**Figure 3**A). Two-rowed barleys from Europe and Turkey formed a large cluster derived from a six-rowed ancestor in Türkiye *c.* 3,500 ya. Two-rowed barleys from Iran, Syria and Iraq were derived from six-rowed ancestors in the same region *c.* 6,000 ya. (**Figure 3**A). The transition from two-rowed to six-rowed type was identified on two occasions, with one in Türkiye at *c.* 3,000 ya, and the other in Georgia at c. 2,500 ya. (**Figure 3**A). Phylogenetic analysis with the *vrs*1 gene haplotypes revealed a similar pattern (**Figure 3**B). Haplotypes corresponding to the two-rowed type were derived from those potentially producing the six-rowed type (**Figure 3**B).

Recent research proposed that early agriculture developed in diverse ways over the Fertile Crescent zone from the southern Levant to the Zagros, and to Central Anatolia (Riehl, et al. 2013; Arranz-Otaegui, et al. 2016; Baird, et al. 2018). We thus sieved the literature for early archaeological sites in the Fertile Crescent zone from the southern Levant to the Zagros and to Central Anatolia for records identifiable as domesticated barley (with non-brittle rachis). We identified 18 sites that recorded remains of domesticated barley and dated earlier than 7,000 cal. B.P. These sites formed three clusters (**Figure 4**). The region, stretching from Dja’de el-Mughara in Northern Syria to Israel along the Jordan Rift Valley to Beidha Jordan, has dense earlier archaeological sites recorded remains of domesticated barley, dated back to as early as 8,200 cal. B.P. at Tell Aswad AW2 in Syria, while semi-domesticated types (with brittle rachis but larger grains) present in site Mureybet III (9,500 cal. B.P.) and Dja’de el-Mughara (9,310 cal. B.P.) in Syria. Four sites in the Foothills of the Zagros Mountains recorded domesticated barley 7,940 cal. B.P. in Jarmo and Tepe Sarab. Southern Anatolian Plateau in Türkiye has four archaeological sites that recorded remains of domesticated barley, dated to 8,014–7,935 cal. B.P. at Asikli Höyük. Across the three clusters of archaeological sites, all the earliest records of domesticated barley were two-rowed types with hulled grains. Six-rowed type and hull-less grain occurred in the archaeological record at a later stage. In the Fertile Crescent along Syria, Israel and Jordan, the earliest six-rowed type was recorded at Ghoraifé Phase II and Ras Shamra, 7,500 cal. BC. The six-rowed type appeared at a similar time in the Southern Anatolian Plateau at 7,650 cal. B.P. at Canhasan III, and in the Foothills of the Zagros Mountains, at 7,420 cal. BC at Khuzistan (**Table 1**). Earlier records of barley remains were reported from sites in the Jordan Rift Valley dated to 23,000 to 21,000 yBP but were believed to be wild types (Kislev, et al. 1992; Snir, et al. 2015) (**Table 1**).

**Figure 4.**
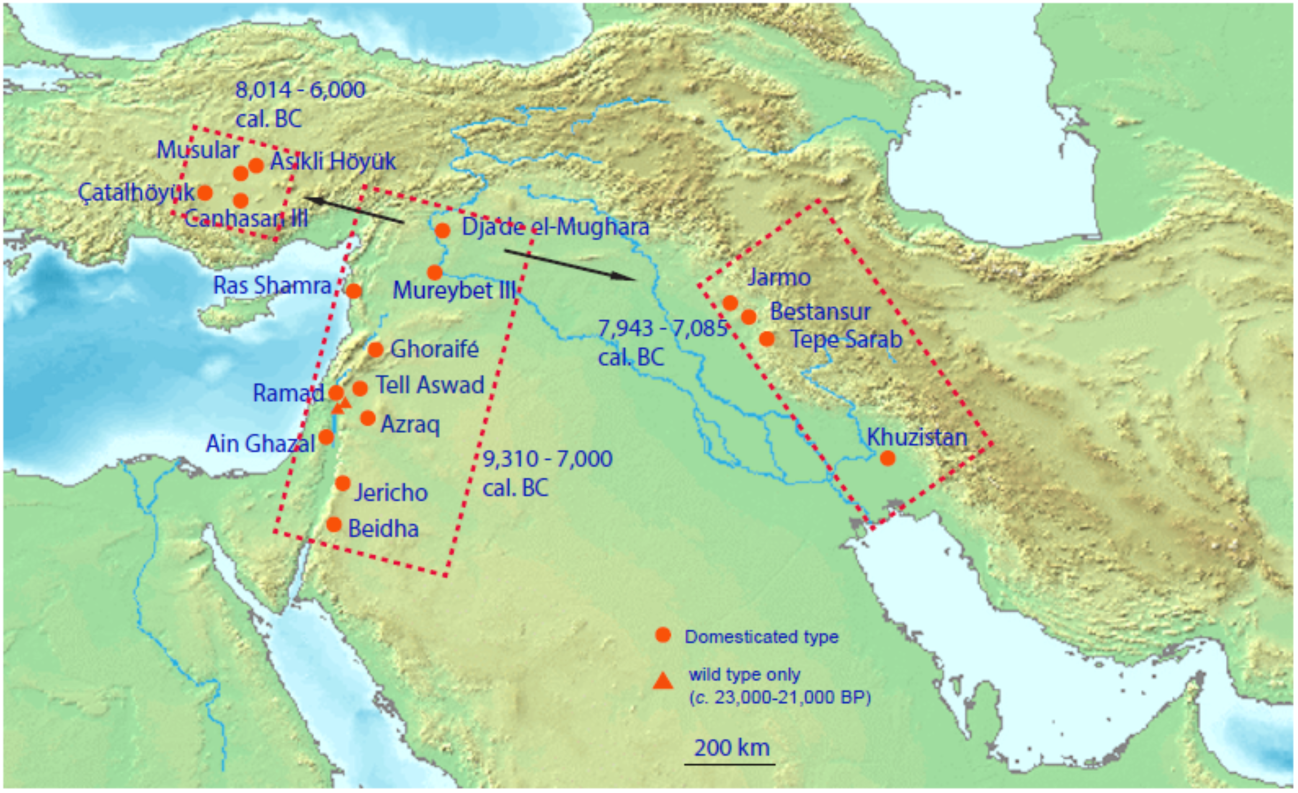
Map and distribution of the archaeological sites. It showing the archaeological sites in the Fertile Crescent from the southern Levant to the Zagros and Central Anatolia with recorded remains of domesticated barley. The base map was adopted from Google Earth.

**Table 1.**
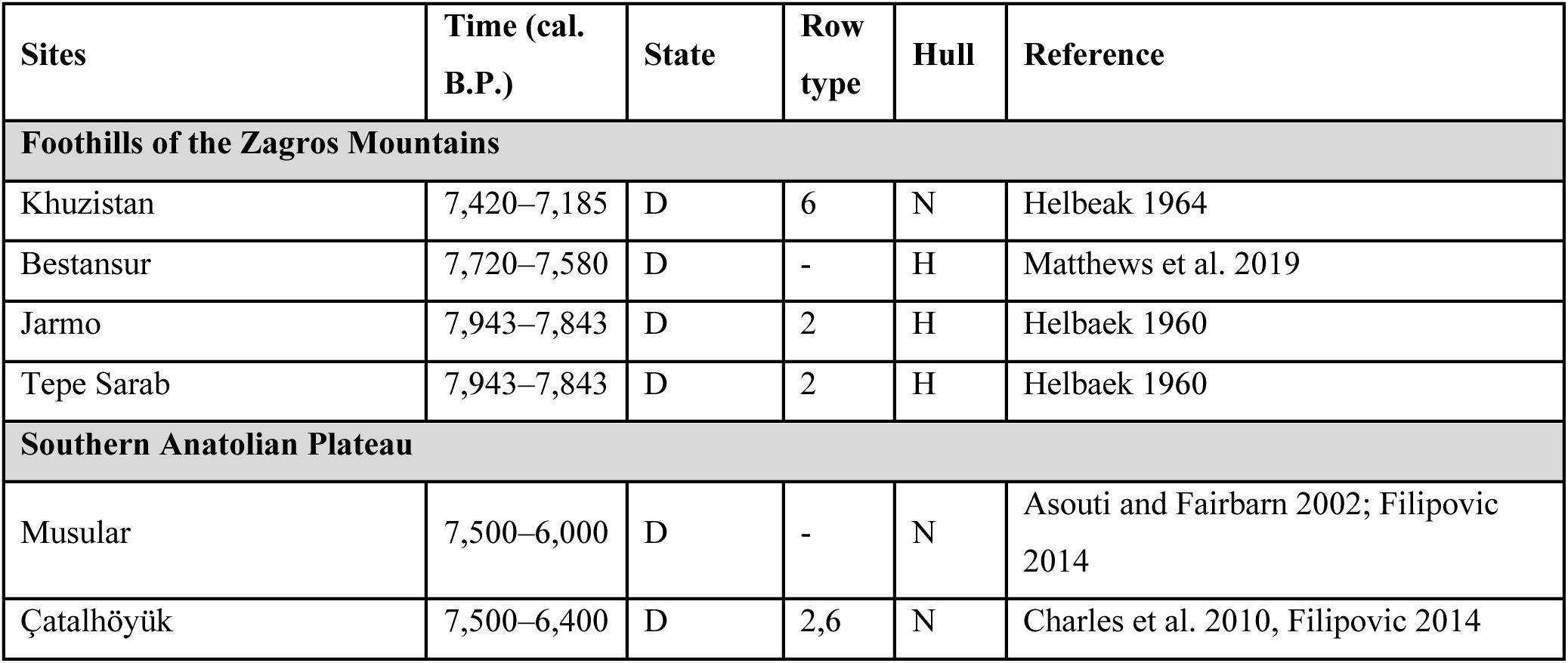
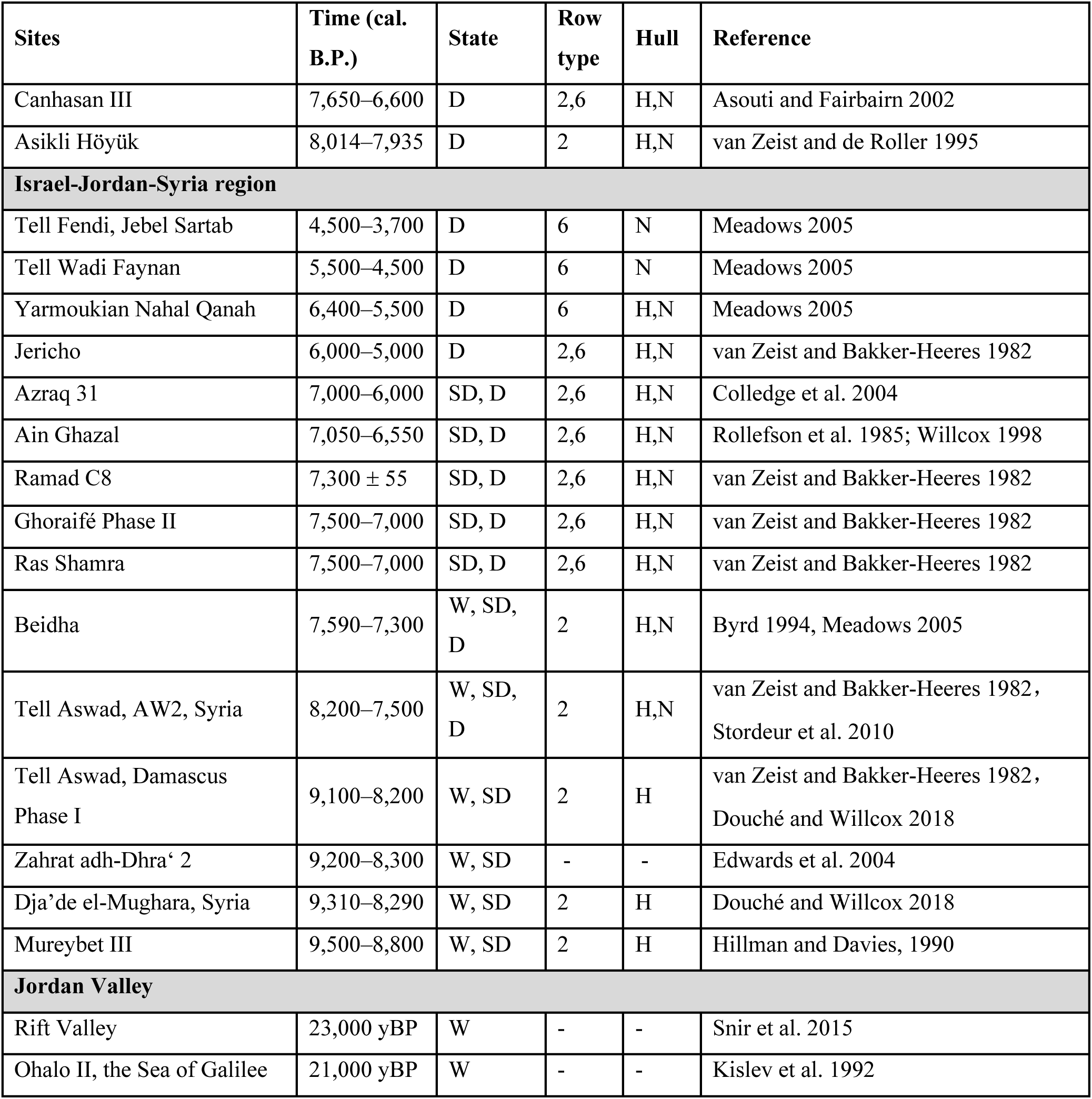
Information of early archaeological sites. Early archaeological sites with barley remain in the Foothills of the Zagros Mountains, Southern Anatolian Plateau, Israel-Jordan-Syria region, and Jordan Valley. D: domesticated form (with non-brittle rachis); SD: semi-domesticated form (with brittle rachis but larger grains); W: wild forms (with brittle rachis and generally small grains). H: hulled grain; N: hull-less grain.

Despite the earliest records of domesticated barley being two-rowed types across the three regions, records of two-rowed types in archaeological sites became infrequent since *c.* 9,000 B.P. and completely absent from archaeological records since c. 6,000 B.P. until c. 3,000 B.P. (**Figure 5A**). Meanwhile, six-rowed barley dominated archaeological records since its later occurrence at c. 10,500 B.P. and continuously to be till recent (**Figure 5A**). The rapid transition from two-rowed barley to six-rowed barley only several hundred years since it was fully domesticated may be related to the climate change in the region since c. 11,000 BP ago with a significant increase of rainfall in the region (**Figure** 5B). Early farmers would choose and rapidly adopt the six-rowed type as the six-rowed type may produce three-fold higher yield than the two-row types in the environment with sufficient water supply.

**Figure 5.**
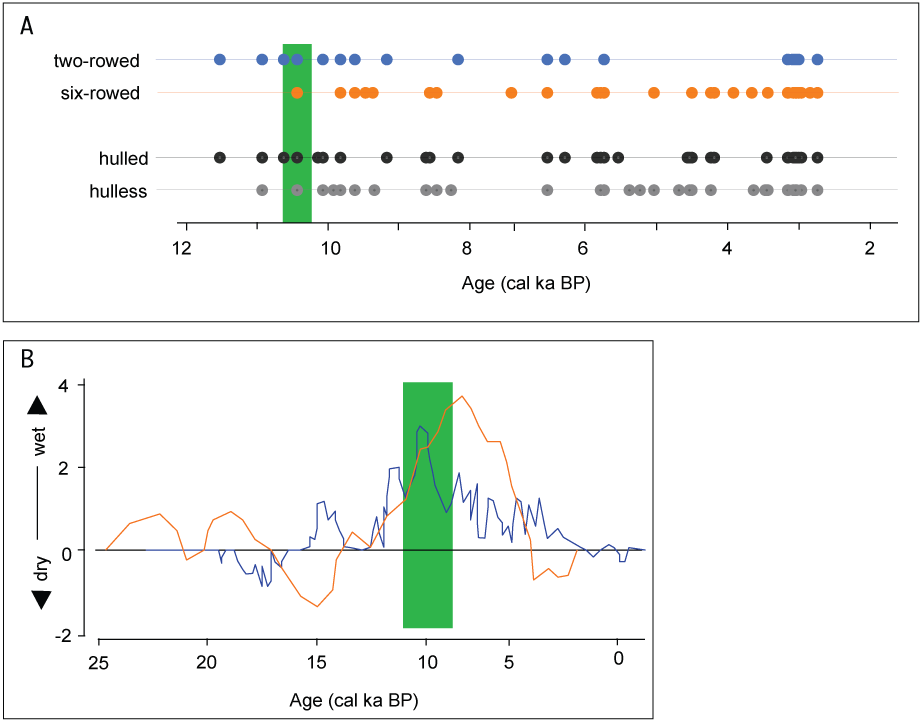
Pattern of barley row-type and reconstructed rainfall changes in archaeological records. A, Pattern of two-rowed barley and six-rowed barley in the archaeological records; **B,** Reconstructed rainfall changes during the past 25,000 years in the Israel-Jordan-Syria region. The blue line was from Grant, et al. (2016), and the red line was from Grant, et al. (2017).

## Discussion

### Early two-rowed barley is extinct, and modern tow-rowed types were derived from domesticated six-rowed, rather than a second and independent domestication

Archaeological records indisputably revealed that the earliest domesticated barley was the two-rowed type (Helbaek 1970; Hillman and Davies 1990; Douché and Willcox 2018). Archaeologists have long observed that the two-rowed types were replaced by the six-rowed shortly after becoming domesticated (Helbaek 1961; Rollefson and Simmons 1985; Willcox 1998). Unmistakable remains of nonbrittle two-rowed barley (an indication of domestication) came from phase II (ca. 10,200–9,550 cal. BP) in Tell Aswad (Tanno and Willcox 2006) and from ca. 9,450 to 9,300 cal. BP Jarmo, Iraq(Helbaek 1970; Morabito, et al. 2022). Given the distinct genetic and phytogeographic pattern of two-rowed barleys and six-rowed barleys, it has been argued that the two types have followed entirely independent evolutionary paths that have been genetically and geographically distinct from the beginning of their history(Takahashi 1955; Takahashi and Hayashi 1964). Our maximum likelihood phylogenetic reconstruction showed that the common ancestor of modern barley was six-rowed, and molecular dating suggested that all modern two-rowed barley originated from the six-rowed type not earlier than 3,500 BP. Genetic data reflect phylogenetic history and population genetics processes, such as selection and extinction (Allaby and Brown 2003). The most plausible explanation is that the original domesticated two-rowed type may be extinct. The modern two-rowed barleys were likely derived from six-rowed domesticated barley, not stemming from the original and early form of two-rowed domesticated barley in the Fertile Crescent, nor a second and independent introduction of wild barley into cultivation.

Thus, the two-rowed types in modern domesticated barley, coinciding with distinct non-brittle rachis haplotypes, are likely the results of two chronological evolutionary events several thousands of years apart, other than two entirely independent evolutionary lines. The change of row number, either in the earlier transition from two-rowed to six-rowed or the late reappearance of two-rowed, was likely through mutations. Row number changes in experimental material due to mutagenic agents have been observed(Ehrenberg, et al. 1961). Recently, Casas, et al. (2018) identified a two-rowed revertant through a mutation in the *vrs1* gene in a Spanish six-rowed population. This proved that the reverse transition from six-rowed to two-rowed type was straightforward and feasible. It is worth noting that six-rowed barley with Type W non-brittle rachis has been observed in regions of northern Afghanistan and Iran (Takahashi, et al. 1963), and Turkey (Takahashi and Hayashi 1964), implying that only one step of mutation would be required for the transition from six-rowed to Type W two-rowed barley. The re-appearance of two-rowed barley since c. 3,500 years ago in regions such as Tukey and Iran may be related to the drying climate at the time (Grant, et al. 2016; Grant, et al. 2017). Under dry and hot conditions, two-rowed barley could have a yield advantage over the six-rowed barley, thus being selected. However, for this hypothesis to withstand further scrutiny, a demonstrated molecular mechanism of transition from six-rowed to two-rowed type is crucial in future research.

### Monophyletic origin of modern domesticated barley and reconciliation with the genetic and geobotanic pattern

Using two independent and large datasets and dated phylogenetic construction, we demonstrated that modern domesticated barley has a monophyletic origin. The primary challenge to the hypothesis of the monophyletic origin of barley was its incompetence in satisfactory explanation of distinct genetic and phytogeographic patterns of barley distribution (Clark 1967). It has long been observed that the two-rowed barleys from the West (the occidental region of the Old World) were of the btr1btr1/Btr2Btr2 non-brittle rachis type (Type W), whereas six-rowed barleys from the East (the oriental region) were predominantly Btr1Btr1/btr2btr2 type (Type E) (Takahashi 1955; Zohary 1959; Clark 1967), which was explained as the result of a diphyletic origin (Takahashi 1955; Takahashi, et al. 1963) before the discovery of the common and ancient origin of hull-less barley (Komatsuda, et al. 2007). Our findings of the two row types, coinciding with distinct non-brittle rachis haplotypes, resulting from two chronological evolutionary events several thousands of years apart, provide a reconcilable framework for explaining domesticated barley’s genetic and phytogeographic pattern. Although our current analysis could not identify the timing of the evolutionary transition of the btr1 (Type W) and btr2 (Type E) non-brittle rachis haplotypes type, Pourkheirandish, et al. (2018) confirmed that the btr1- and btr2-type barleys emerged at different times and observed that the sequence of btr2-type haplotypes are closer to wild barley than the btr1-type haplotypes. The ancient two-rowed domesticated barleys and their directly derived six-rowed thus likely emerged as btr2 non-brittle rachis haplotype, whereas btr1-type six-rowed barley entered cultivation at a later stage before the re-appearance of two-rowed barley. The precise genetic mechanism of the transition of non-brittle rachis type warrants further study. The transition, however, is likely the result of neutral genetic mutations other than through natural selection.

Archaeological evidence suggested that both hulled and hull-less barley (presumably six-rowed and with Type E non-brittle rachis) was cultivated in northern Europe in pre-historical periods (Bogaard and Jones 2007), but hull-less barley largely disappeared from the archaeobotanical record from the Iron Age/Roman periods in Europe (Lister, et al. 2018), while two-rowed barley (Type W) becomes important in Europe since Roman and medieval periods, coinciding with a transition from predominantly being food to being brewed for beer since the Iron/Roman period. Thus, the predominantly presence of btr1 type non-brittle rachis in Europe is likely the result of the westwards spreading of the re-appeared two-rowed barley with btr1 type non-brittle rachis from regions such as Türkiye from the Iron Age/Roman periods.

### Barley expanding from the west Fertile Crescent

Previous research on haplotype distribution proposed that early barley domestication in the Zagros and further east in Iran was independent of its domestication in the Western Fertile Crescent (Morrell and Clegg 2007; Saisho and Purugganan 2007). Our analysis of global barley landraces and wild accessions throughout the Fertile Crescent and beyond revealed that modern barley is phylogenetically related to wild barley from Israel and Jordan, as Mascher, et al. (2016) have previously observed. Thus, barley was most likely first domesticated in the Western Fertile Crescent and spread to the Fertile Crescent’s eastern area in the Zagros Mountains’ foothills and westwards to southeastern Anatolia over 1,000 to 2,000 years. Outside the Western Fertile Crescent, archaeological records of domesticated barley in the Fertile Crescent’s eastern area in the Zagros Mountains’ foothills dated back to 7,080 cal. B.C. at Jarmo (Helbaek 1964; Riehl, et al. 2013), and in southeastern Anatolia, approximately 7,800 B.C. (Van Zeist and de Roller 1994; van Zeist and de Roller 1995), forming the basis for the hypothesis of the polyphyletic origin of barley domestication. However, unmistakable remains of domesticated barley with non-brittle rachis from phase in Tell Aswad dated to c. 9,100–8,200 cal BC (Van Zeist and Bakker-Heeres 1982), and 9,500–8,800 in Mureybet II (Hillman and Davies 1990), which is 1,000-2,000 years older than those from Fertile Crescent’s eastern area and Anatolia. Likely, barley domestication and cultivation were 1,000-2,000 years earlier in the Western Fertile Crescent than in the east area in the Zagros Mountains and southeastern Anatolia. Further, archaeological evidence also suggests that the use of wild barley, e.g. harvesting grains from the wild or even cultivating the wild barley, may have occurred much earlier in the Jordan Rift Valley in the Western Fertile Crescent dated to 23,000 to 21,000 yBP, 10,000 years before morphological domestication (Kislev, et al. 1992; Snir, et al. 2015). Agriculture began in the Fertile Crescent of Southwest Asia 10,000 to 9,000 BC and spread subsequently across western Eurasia, reaching central Anatolia by c. 8,300 BC with human migration and exchange of farming culture and technologies (Baird, et al. 2018; Feldman, et al. 2019). Recently, research of ancient human genomics suggested a common origin of human populations in regions containing modern-day Iran, Iraq, Israel, Palestine, Jordan, Lebanon, Syria, and Türkiye before the westward spread of agriculture in the Fertile Crescent area (Kilinç, et al. 2016). Early humans may have moved from the West Fertile Crescent westwards to Anatolia and south-eastwards’ to the eastern area in the Zagros Mountains and beyond (Marchi, et al. 2022). Together with a pulse of human migration, archaeological evidence also showed the movement of crops between settlements and regions in the early phases of the Neolithic c. 10,000 BC (Arranz-Otaegui, et al. 2016; Baird, et al. 2018). Thus, barley domestication and cultivation were first developed in the West Fertile Crescent. The exchange of crop materials and farming technologies led to the spread of farming within the Fertile Crescent and eventually to the Zagros and Central Anatolia (Arranz-Otaegui, et al. 2016) within 1,000-2,000 years.

It is not disputed that the six-rowed barley replaced the two-rowed types shortly after becoming domesticated. The transition from the two-rowed type to the six-rowed type might be related to the change in climate at the time. Indeed, Ŝeveluha (1957) reported that changes in the environment of the growing barley plants had induced the mutation from the two-rowed to the six-rowed state (Clark 1967), Helbaek (1970) thus proposed that the mutations creating the six-rowed barley from the two-rowed may have coincided with the first use of irrigation. It has long been noted that irrigation increases grain width for any barley variety (Harlan 1914). Six-rowed types bore an obvious advantage for early farmers as they would produce three folders of grains compared to the two-rowed types in environments with sufficient water supply. Apart from the possible irrigation in early agricultural practice, increased rainfall would significantly differentiate the yield potential of two-rowed and six-rowed barley in the field (del Moral, et al. 2003). The rapid transition from two-rowed to six-rowed barley may be related to climate change in the region, with a significant increase of rainfall in the region of 1,1000 – 9,000 yBP. Early farmers would have consciously chosen and rapidly adopted the six-rowed types, given their greater yield over the two-rowed. Thus, the rise of six-row types in early agriculture may be explained by its higher fecundity potential, derived from changing to favourable climate and advanced farming practices.

### Conclusions

The origin of domesticated barley has been debated without a clear consensus over the past century. A polyphyletic origin has gained some acceptance based on botanical and comparative morphological evidence (Schiemann 1948)and, recently, genetic research (Morrell and Clegg 2007; Poets, et al. 2015; Civáñ and Brown 2017; Pankin, et al. 2018). Using two independent datasets with a large number of samples and genome-wide genetic markers, we unequivocally demonstrated that modern domesticated barley has a monophyletic origin c. 13,000 BP, consistent with the fact that a single genetic locus controls the presence of hull-less grains in modern domesticated barley (Taketa, et al. 2008). Together with phylogenetic reconstruction, a thorough examination of current archaeological records suggested the Israel-Jordan area in the west Fertile Crescent was most likely the region where barley was first domesticated, and the domesticated crop then spread to the east area of Fertile Crescent and westwards to east Anatolia in 1,000-2,000 years as the results of pulses of human migration from the Fertile Crescent and the exchange of farming practices and technologies. The original and early domesticated two-rowed barley may be extinct. The modern two-rowed barleys were likely derived from six-rowed domesticated barley, not earlier than 3,500 years ago in modern-day Türkiye and Iran. We suggest that the two distinct row types are likely the results of two chronological events thousands of years apart along the same evolutionary line. Our results provide a reconcilable and testable framework for explaining domesticated barley’s genetic and phytogeographic pattern (**Figure 6**) and pave the way for a consensus on the origin of barley domestication.

**Figure 6.**
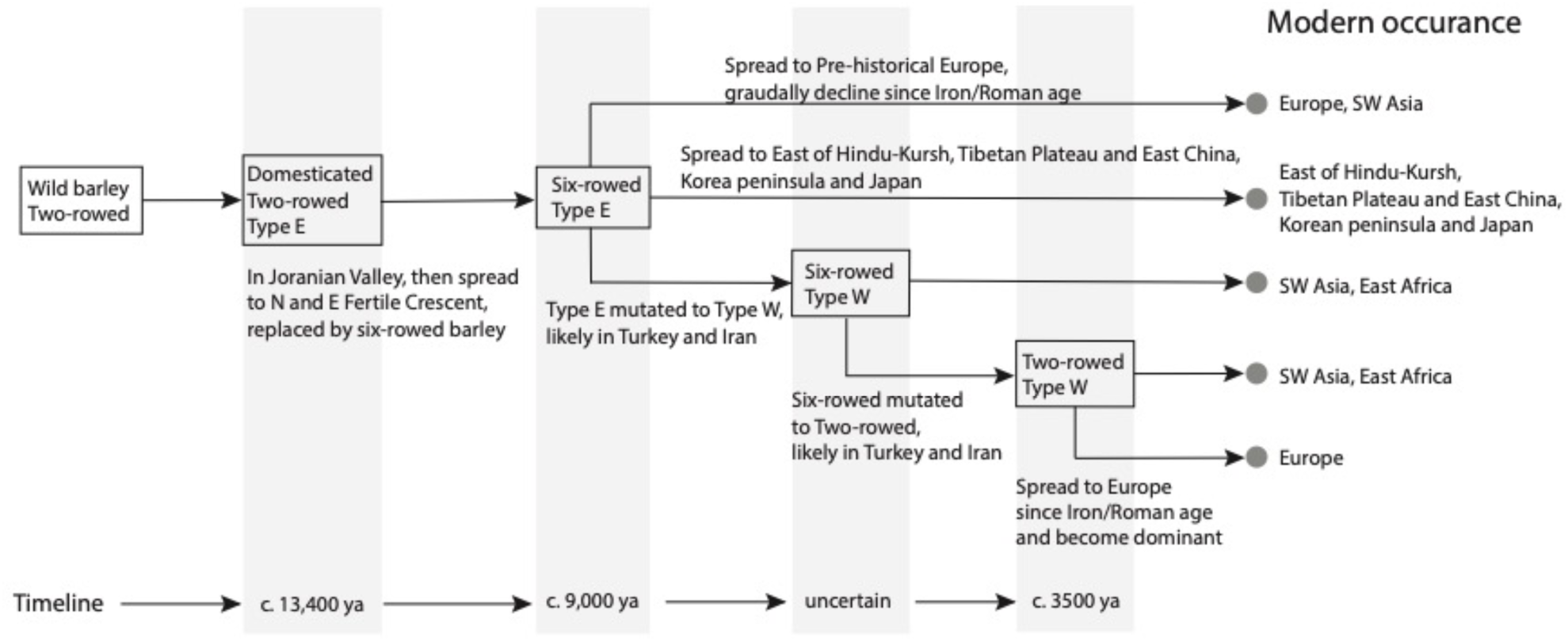
**The evolutionary pathway of domesticated barley under the framework of monophyletic origin.**

## Acknowledgement

We greatly thank the help from members of the Western Crop Genetics Alliance (WCGA, Australia) and colleagues from the International Barley Hub (IBH, UK). We also sincerely thank the support provided by China National GeneBank (CNGB).

## Figure and Table legends

**Supplementary Table 1.** List of 340 high-coverage sequenced barley accessions.

**Supplementary Table 2.** List of 5,448 low-coverage sequenced barley accessions

**Supplementary Table 3.** Abbreviations for countries and regions

